# Developmental delay in attaining adult levels of motor excitability in children and adolescents with Tourette syndrome: a mega-analysis study

**DOI:** 10.64898/2026.05.14.724875

**Authors:** Stephen R. Jackson, Valerie Brandt, Christine A. Conelea, Kevin J. Black, Donald L. Gilbert, John Piacentini, John Rothwell, Yulia Worbe, Katherine Dyke

**Affiliations:** School of Psychology, University of Nottingham, UK; University of Nottingham Centre for Neuromodulation, Neurotechnology & Neurotherapeutics, UK; Department of Psychology, Centre for Innovation in Mental Health, University of Southampton, UK; Clinic of Psychiatry, Social Psychiatry and Psychotherapy, Hannover Medical School, Hannover, Germany; Department of Psychiatry & Behavioral Sciences, Masonic Institute for the Developing Brain, University of Minnesota, Minneapolis, MN, USA; Departments of Psychiatry, Neurology, Radiology, and Neuroscience, Washington University in St. Louis, St. Louis, MO, USA; Cincinnati Children’s Hospital Medical Center, Cincinnati, OH, USA; UCLA Semel Institute for Neuroscience and Human Behavior, UCLA, Los Angeles, CA, USA; UCL Queen Square Institute of Neurology, University College London, UK; Paris Brain Institute - ICM, Sorbonne University, INSERM, CNRS Paris, France; Department of Clinical Neurophysiology, Assistance Publique des Hôpitaux de Paris, Hôpital Saint Antoine, 75012, Paris, France

**Keywords:** Tourette syndrome, Motor excitability, Transcranial magnetic stimulation (TMS), Motor threshold, children and adolescents

## Abstract

Tourette syndrome (TS) is a neurodevelopmental disorder of childhood onset characterised by vocal and motor tics and is associated with cortical-striatal-thalamic-cortical circuit [CSTC] dysfunction. TS often follows a developmental time course in which tics become increasingly more controlled during adolescence. However, many individuals continue to have debilitating tics into adulthood. This indicates that there may be important differences between adults with TS for whom the clinical phenotype is more stable, and children and adolescents with the disorder who may be undergoing developmental neuroplastic changes linked to the reduction of their tics.

Previous studies have used transcranial magnetic stimulation (TMS) to investigate changes in cortical motor excitability in individuals with TS, including measurement of resting motor threshold (RMT). However, the findings from these studies have been mixed, have varied between adult and child samples, and have often been based on small sample sizes. Here we report a multi-centre, mega-analytic, study in which RMT data collected from children and adults with TS at multiple research centres was pooled for analysis.

Results confirmed that mean RMT was significantly increased in individuals with TS compared to neurotypical controls. However, this result can be explained by the more important findings that: (a) RMT for adults with TS did not differ from that of neurotypical adults; and (b) the rate that RMT *decreases with age* during childhood and adolescence is reduced in individuals with TS compared to controls. Thus, while neurotypical individuals reach an adult RMT level by ~12-13 years of age, individuals with TS are substantially delayed in doing so, and do not reach an adult RMT level until much later, at ~24 years of age.

We conclude therefore that differences in measures of cortical excitability between children and adolescents with TS and chronologically age-matched neurotypical controls may likely reflect a developmental delay in the maturation of functional brain networks in individuals with TS, which may normalise with age.

## Introduction

Tourette syndrome (TS) is a neurodevelopmental disorder of childhood onset that is characterised by the presence of chronic vocal and motor tics [1]. Tics are involuntary, repetitive, stereotyped behaviours that occur with a limited duration, often many times in a single day [1]. People with chronic motor tics but no phonic tics have similar clinical features, family history, course, and treatment response, and hereinafter we will not distinguish these patients from those with TS.

TS often first presents during early childhood (~4-7 years). Tic severity is frequently maximal between 11-14 years, with tics decreasing by early adulthood for about 2/3 of individuals [1]. Thus, TS typically follows a developmental time course in which the tics reduce in frequency and intensity across adolescence. However, a substantial minority of individuals continue to have debilitating tics into adulthood, with symptoms becoming more severe in some cases and resistant to treatment [1].

While the neurobiological basis of TS remains unclear, it is generally acknowledged that cortical-striatal-thalamic-cortical circuits (CSTC) are dysfunctional in TS. Specifically, it is suggested that striatal disinhibition, in which subsets of striatal projection neurons becoming active within inappropriate contexts, results in the disinhibition of thalamo-cortical projections [2] and hyper-excitability of cortical motor regions [2–5], which in turn leads to the occurrence of tics [2].

Alterations in cortical excitability in TS have been studied using Transcranial Magnetic Stimulation (TMS) [3–5]. TMS can be used to stimulate the primary motor cortex and induce a motor evoked potential (MEP) in a targeted muscle; it is therefore a useful tool to measure corticospinal excitability (CSE) non-invasively at rest or during the preparation and execution of movement. A fundamental property quantifiable by TMS is the measurement of an individual’s motor threshold. Motor threshold is defined as the minimum intensity of stimulation required to reliably induce an MEP of a given amplitude in a target muscle; either at rest (termed resting motor threshold - RMT), or when the muscle is partly activated (termed active motor threshold - AMT).

A key theoretical construct is the ‘gain’ in CSE. This can be defined as the rate at which CSE increases with increased inputs (e.g. increasing TMS pulse intensity). This construct can be operationalised in several ways but is most often measured as the gradient (slope) of the TMS recruitment curve. TMS recruitment curves are assessed by using a stimulus-response TMS technique, where the intensity of TMS is systematically increased from RMT to measure the intrinsic capability of the motor cortex to ramp up excitability in the resting muscle (i.e., the ‘gain’ in motor excitability). The gain of CSE can also be measured during the preparation of a volitional movement, where the TMS intensity is not altered, but the time at which the TMS pulse is delivered, relative to the onset of movement, is varied systematically. Typically, the closer to the onset of movement that TMS is delivered, the greater the MEP amplitude [6]. In this way the gain in motor excitability can be quantified from the gradient of MEPs obtained during movement preparation.

Several key findings are relevant to the current study: First, several studies have reported previously that RMT in individuals with TS does not differ from that of age matched controls [5, 7–13], whereas others have reported that RMT is significantly increased in individuals with TS [14, 15]. That is, *increased* TMS stimulation is required to produce an MEP of a given amplitude in individuals with TS compared to that required to produce an equivalently sized MEP in neurotypical individuals. Importantly, it is suggested that while equivalent RMTs between TS patients and controls indicate that the neural populations recruited by TMS at threshold were similar, RMT differences might indicate that different neural populations are recruited, leading to different levels of motor excitability [15]. Second, a number of studies have demonstrated that the gain of motor cortical excitability is reduced in individuals with TS. This is the case for both TMS-induced increases in motor excitability (i.e., TMS recruitment curves) [4, 14, 16] and for gains in motor excitability observed during motor preparation, i.e., preceding the execution of a volitional movement [5, 14, 17]. Importantly, the gain in cortical excitability is thought to depend upon the distribution of excitability within the population of corticospinal neurons (i.e., recruitment of neurons with different levels of excitability): thus, a shallower gain function in TS reflects a wider spread of excitability within this population [15].

Unfortunately, drawing conclusions from existing literature is made difficult by several factors. First, many early studies were based on small samples that likely lacked sufficient statistical power to detect between group differences in RMT. A recent meta-analysis of RMT values reported in such studies confirmed that with sufficient statistical power, RMT is significantly increased (standardized mean difference = 0.85) in individuals with TS compared to age-matched neurotypical controls [18]. Second, many studies investigating cortical excitability in TS using TMS have been conducted in *adults* with TS, for whom the clinical phenotype is likely more stable than children and adolescents with the disorder. Consequently, the results may not generalize across the developmental course of TS. Third, in neurotypical subjects, the motor threshold has been shown to *decrease* with age during childhood and adolescence with substantial change occurring before mid-adolescence [19, 20]. This is important as many of those studies that have investigated RMT in children and adolescents have used small samples with a restricted age range.

The current study aimed to examine the developmental trajectory of RMT values in TS by conducting a mega-analysis of cross-sectional RMT data from a sufficiently large, pooled sample of children, adolescents, and adults with TS and age matched neurotypical individuals. Our aims were as follows: 1- to assess group level differences in RMT to address previous discrepancies within previous literature. 2- to explore predictors of RMT including age, sex at birth, and tic severity. 3- to quantify how RMT changes with age and to evaluate if this differed as a function of diagnostic group.

## Methods

### Sample

As part of a Medical Research Council (UK) Partnership grant [MR/Z505055/1], data from 426 participants were pooled from eleven studies completed by the participating research centres. The combined sample consisted of 235 individuals with TS or chronic tic disorder (67 female at birth, 28.5%, mean age = 20 ± 9.21 years, age range: 8.0 – 65 years) and 191 neurotypical controls (60 female at birth, 31.3%, mean age = 21 ± 9.8 years, age range: 8 – 66 years). Care was taken to ensure that individuals only contributed one data set in this analysis. Consequently, the total numbers included and reflected in Table 1 may be lower than in original publications as participants who took part in several studies will only have had one entry included. All sites contributing data to this analysis are established TS research centers with expertise in recruitment and assessment of tics.

**Table 1:** Key study aspects for the 11 contributing studies to the pooled data set. MP= monophasic pulse, BP= biphasic pulse. For neuronavigation template indicated use of a standard scaled template e.g. MNI atlas, where as anatomical indicated use of individual RI scans. Note several studies are ongoing and/or in prep and therefore do not have available citations.

| Study number & reference | Target muscle | TMS and coil | Neuro navigation | TS (N) | CS (N) | Age range (years) |
| --- | --- | --- | --- | --- | --- | --- |
| 1: Dyke et al, 2019.[23] | Right FDI | Magstim 200 (MP)<br>Figure 8, 70mm | Y Brainsight (template) | 7 | 0 | 16:33 |
| 2: N/A | Right FDI | Magstim 200 (MP)<br>Figure 8, 70mm | Y Brainsight (template) | 8 | 7 | 10:24 |
| 3: N/A | Right FDI | Magstim 200 (MP)<br>Figure 8, 70mm | Y Brainsight (template) | 9 | 11 | 14:21 |
| 4. Sigurdsson et al, 2020 [24] | Right FDI | Magstim 200 (MP)<br>Figure 8, 70mm | Y Brainsight (Anatomical/template) | 14 | 10 | 12:35 |
| 5. Batschelett et al, 2023[25] | Right FDI | Magstim 200 (MP) Circular, 90mm | N | 27 | 29 | 8:12 |
| 6. Brandt, et al, 2014 [12] | Right APB | Magstim 200 (MP) Figure 8, 70mm | N | 14 | 15 | 18:39 |
| 7. N/A | Right FDI | Magstim 200 (MP) Figure 8, 70mm | Y Brainsight (template) | 17 | 17 | 19:54 |
| 8. Conelea et al, 2023 [26] | Right APB | Magstim Super Rapid2 (BP) Figure 8, 70mm air-cooled | Y Brainsight (anatomical) | 48 | 0 | 12:21 |
| 9. N/A | Right FDI | Magstim 200 (MP) Figure 8, 70mm | Y Brainsight (template) | 16 | 27 | 17:65 |
| 10. Tubing et al, 2018 [27] | Right FDI | Magstim 200 (MP) Figure 8, 50mm | Y Brainsight (anatomical) | 14 | 22 | 20:51 |
| 11. N/A | Right FDI | Magstim 200 (MP) Figure 8, 70mm | Y Brainsight (template) | 61 | 53 | 14:19 |

All individuals with TS or chronic tic disorder were assessed by trained experts using the Yale Global Tic Severity Scale (YGTSS) [21]. This validated clinical rating scale assesses symptoms occurring in the previous week including tic type/location, frequency, intensity, complexity and interference. The subscales for total motor and total phonic scores (each scored 0-25) are used in this analysis.

All studies in this analysis obtained ethical approvals through their respective ethics committees.

### Measurement of resting motor threshold

All studies followed the well-established guidelines and protocols for calculating resting motor threshold [22] which are summarised as follows. **1: Identification of the motor ‘hotspot’ for a target muscle at rest**. This is achieved by systematically moving the coil in small increments around the target region, delivering single TMS pulses, and measuring the amplitude of motor evoked potentials (MEPs) in the target muscle. The area which consistently yields the largest MEPs in the target muscle is deemed the ‘motor hotspot’ and is tracked throughout the experiment. All studies included in this analysis applied TMS to the left hemisphere, targeting either the first dorsal interosseous (FDI) or abductor pollicis brevis (APB) muscle in the right hand. The majority of studies (exception of studies 5 and 6) used neuro navigational systems to aid coil placement (see Table 1 for study-specific summary). **2. Identification of RMT intensity**. Following the identification of the hotspot the coil is placed over this area, and the intensity of the TMS pulse is gradually decreased until MEPs larger than a fixed intensity (>50μV) are seen approximately 50% of the time in a consecutive train of pulses (10-20). The %maximum stimulator output that archives this is the individual’s RMT.

### Statistical analysis

The analysis presented below were conducted on the full pooled data set (N=426 participants in total). Additional analysis including only individuals <50 years, and analysis excluding data from study 8 (which uses biphasic pulses and has no control group) can be found in the supplementary material. Importantly, neither of these more conservative analyses changed the findings in any significant way.

To ensure there were no statistically significant differences in age between the two groups, independent t-tests were conducted. Chi-squared analysis was also used to assess any significant differences for sex at birth between the groups.

To address our first aim of comparing group level RMT values between diagnostic groups, an independent samples t-test was used. This univariate approach reflects the approach taken by previous studies to identify any differences in RMT (e.g. [9, 13, 14, 28]) and allows for more direct comparisons between previous work.

To address our second aim of exploring demographic factors predictive of RMT we conducted a stepwise multiple regression, in which the following were entered as predictor variables: Sex at birth, Diagnostic Group, Age, Age x group and Age x Sex interaction terms. A second stepwise multiple regression was conducted for the TS group to explore if YGTSS measures added any predictive power (age, sex at birth, YGTSS motor and YGTSS vocal).

To address our third aim of exploring relationships between diagnostic group and age we then fit the data for each group using an exponential function, providing three parameters of interest for each group: The initial RMT amplitude value (A); the rate at which RMT values decay (k), and RMT value at asymptote (C). We used permutation testing (1000 permutations) to assess the rate of decrease in RMT (k). The point at which the asymptote was reached on the X axis (indicating RMT value) was calculated and a further permutation test (1000 permutations) was conducted to confirm any potential differences between these values between the two diagnostic groups. Lastly, to test whether the RMT values for each group differed once they had reached asymptotic levels, an independent groups t-test was conducted.

All data analysis were conducted in Matlab and SPSS statistics. The code and data associated with this manuscript will be made available upon publication.

## Results

The two groups did not differ significantly in either age (*t*(424) = −1.469, *p* = 0.14, d=−0.143) or sex (Yates-corrected chi-square = 1.112, p = 0.292 Effect size (phi / Cramer’s V) = 0.056. Proportion male: TS group = 0.72, CS group = 0.66).

### RMT differs by diagnostic group

RMT for the TS group was significantly higher than that for the CS group (mean RMT: CS group = 47.7 ± 10.9; TS group = 51.5 ± 12.0 %; *t*(424) = −3.45, *p* < 0.001, *d* =.336).

### Diagnostic group and age predict RMT

The stepwise regression mode was statistically significant (F(2,423), =27.05, R^2^ =.11, p=<.001)^=^. Both age (B=−.361, t=−6.408, p=<.001) and diagnostic group (B=−3.386, t=−3.155, p=.002) were significant predictors.

### YGTSS tic severity scores do not predict RMT

The stepwise regression model was statistically significant (F(1,183), = 27.719, R^2^= .105, p=<.001). Age was the only significant predictor in the model (B = −0.413, *t* = −5.169, p < .001).

Further exploration using Pearsons’s correlation (Figure 1) shows a clear lack of statistically significant relationship between YGTSS motor scores and RMT (r(230)=.075,p=.260). There was also no significant correlation between YGTSS phonic scores, which was true with all data include (r(229)=.123, p=.063) and when scores of 0 were excluded, (r(185)=.113, p=.125).

**Figure 1.**
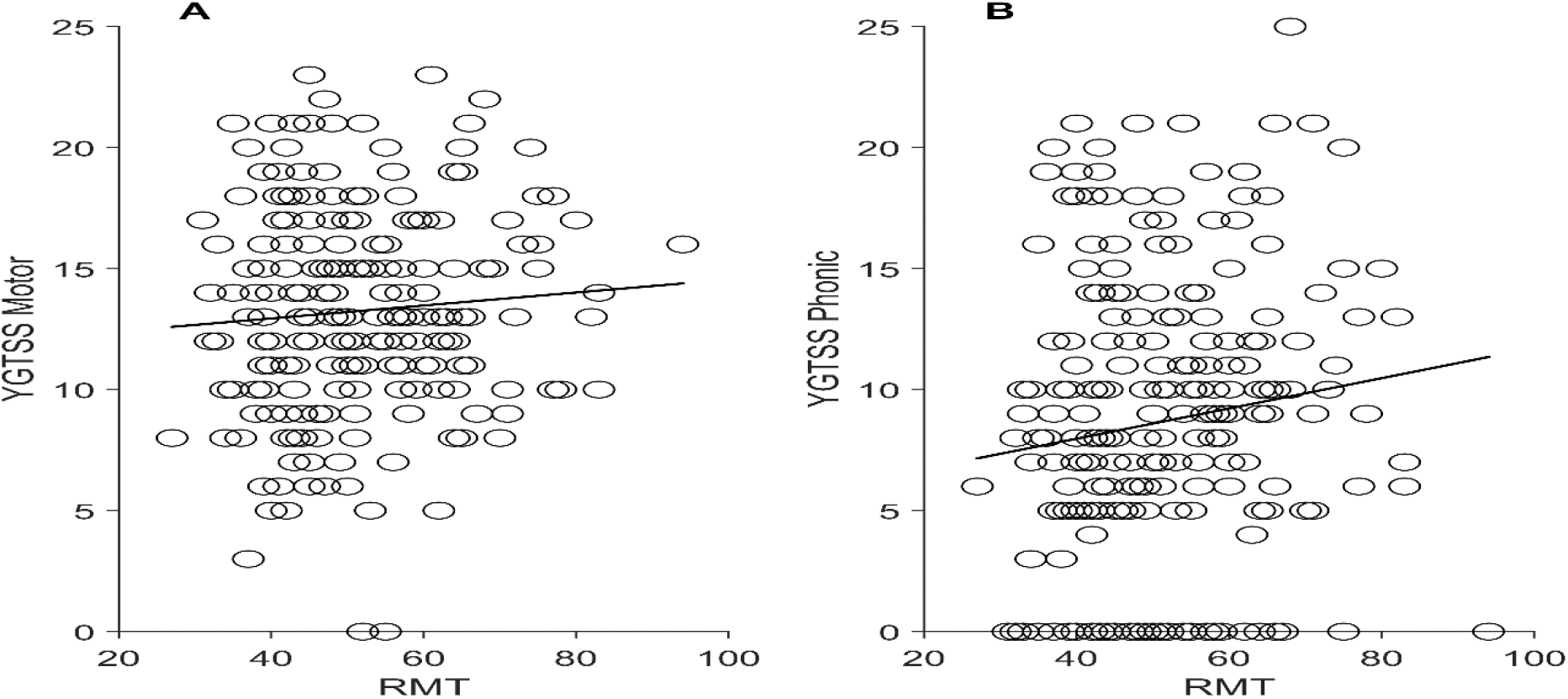
A: YGTSS motor scores plotted against RMT values. B: YGTSS phonic scores plotted against RMT values.

### Differences in RMT by diagnostic group are driven by differing patterns of change in childhood and adolescence

Figure 2 shows a scatter plot of RMT values as a function of age for each group. Inspection of this figure clearly indicates that the RMT data for each group decreases with age but appears to plateau on reaching adulthood. To investigate this further an exponential function was fit to the data from each group. This is shown in equation 1 below.

**Figure 2.**
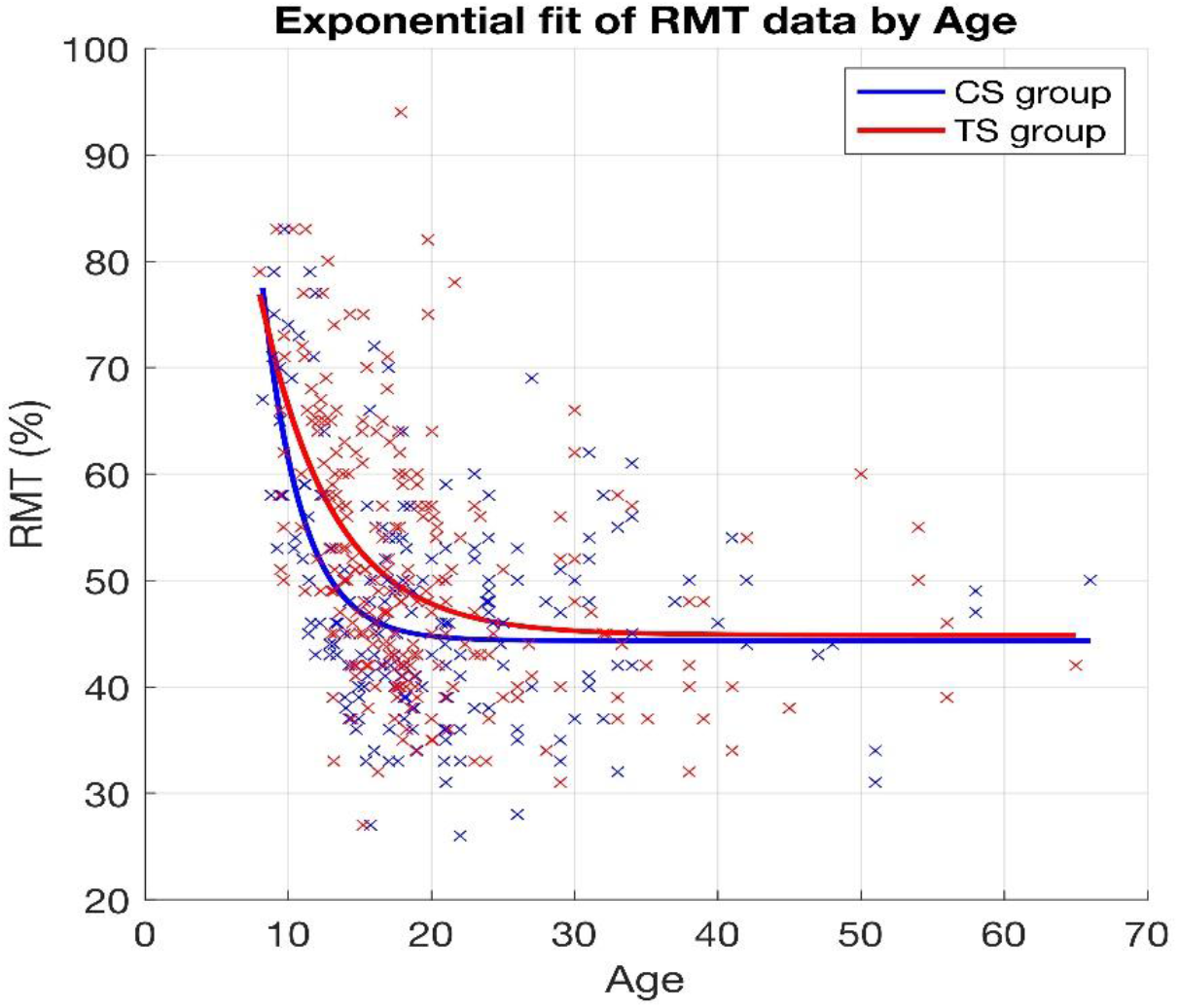
Scatter plot showing individual RMT values as a function of age for the CS group (blue) and the TS group (red). The data from each group was fitted using an exponential function. The solid blue line indicates the fitted function for the CS group and the red line for the TS group. Note both groups show a decrease in RMT with age during adolescence before reaching asymptote as adults.

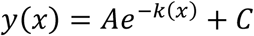

The exponential function produces three parameters of interest: The initial RMT amplitude value (A); the rate at which RMT values decay (k), and RMT value at asymptote (C). The values of these parameters for each group were (CS group: A = 679.4, k = 0.37, C = 44.33; TS group: A = 157.3, k = 0.2, C = 44.86). Inspection of Figure 1 suggests that the rate of decrease in RMT (k) differs both as a function of age and group status. More specifically, the CS group reach adult levels of RMT (C) earlier than the TS group, and that the RMT values at asymptote were similar for each group.

To test for differences in the rate of decrease in RMT (k) during adolescence a permutation test was conducted (10000 permutations). This test revealed that the observed difference in k value between the groups (0.17) was statistically significant (p = 0.007).

The equation below was used to estimate the point at which asymptote was reached for each group. In this case ε = 0.001 (1%).

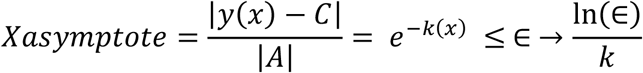

This above equation estimates, for each value of age (x), the distance that value is from asymptote (C). The ratio |y(x) – C| / |A| gives the fraction of the distance that remains. The point at which the x value first reaches asymptote is determined by the value of ε. In this case ε = 0.001 (1%), i.e. we define “reaches asymptote” as “coming within 0.1% of asymptote.” Using this definition, the CS group reached asymptote at 12.51 years of age while the TS group did not reach asymptote until 23.14 years of age. To test this observed difference of 10.64 years, a further permutation test (10000 permutations) was conducted. This test determined that the observed difference in the age at which each group reached asymptotic levels of RMT was significantly different (p = 0.02).

An independent t-test comparing RMT values post asymptote for each diagnostic group did not differ statistically differ (Means: CS group 44.9 ± 8.41, TS group 44.85 ± 8.39; t(206) < 1, p = 0.95, d = 0.01). Suggesting that once RMT had ceased decreasing adult levels of RMT are comparable across groups.

## Discussion

We conducted a mega-analytic analysis of pooled RMT data collected from children and adults with TS and matched neurotypical controls collected from multiple research centres. The results of our study can be summarised as follows. First, consistent with a recent meta-analysis [18] with sufficient statistical power to detect a difference, RMT is significantly increased in individuals with TS compared to age and sex matched neurotypical controls. Secondly, we show that age and diagnostic group significantly predict RMT and that Sex at birth, YGTSS motor, and YGTSS phonic scores do not. Our third finding shows that RMT decreases in childhood/adolescence before reaching a plateau. This rate of change has previously been shown in neurotypical controls [19, 20], however, here we show for the first time that this change may occur more slowly in the TS group. Specifically, we find that RMT reaches asymptotic adult levels much later (~23.1 years) in the TS group compared to age and sex matched neurotypical controls (~12.5 years). This suggests that the maturation of cortical motor excitability is substantially delayed in the TS group.

Notably, once maturation is completed (i.e., asymptotic levels of adult RMT have been reached), our results indicate that RMT values do not differ between individuals with TS and neurotypical individuals. This suggests that any observed difference in RMT between individuals with TS and age-matched neurotypical controls is likely to be explained by an imbalance between samples in the number of individuals who have yet to attain asymptotic levels of adult RMT. This may explain why some, but not all studies have previously reported RMT differences between groups.

These results converge with what is known about the typical developmental time course of TS, in which the tics often reduce in frequency and intensity across adolescence [1]. This symptom-level change has been linked to the delayed maturation behavioural control brain networks that enable top-down control over movement output [29].

Consistent with this proposal, it has been reported that young children and young adolescents with TS exhibit immature functional connectivity within brain networks linked to action control [29, 30]. RMT is also shown to decrease with age during childhood and early adolescence [19, 20], and the rate at which RMT decreases is most likely linked to the rate at which cortical motor networks mature during this period. Our finding that the rate at which RMT decreases during adolescence is significantly reduced in individuals with TS compared to matched controls, leading them to attain adult levels of RMT much later, is consistent with the proposal of a developmental delay in action control networks in TS. It is also noteworthy that our results indicate that RMT is not different in individuals with TS, compared to controls, once they have reached asymptotic adult levels of RMT.

Interestingly, we found no association between clinical measures of motor/phonic tic severity (YGTSS scores) and RMT values, several factors may contribute to this. RMT is a specific measure of primary motor cortex physiology, but overall tic severity likely reflects the summative activity of multiple cortico-striatal networks, including the action control network (responsible for top-down control of motor inhibition) and limbic network (involved in urges to tic and emotion-related impacts on motor output and control). Measurement issues specific to the YGTSS may also play a role here. The YGTSS is a global rating of tic severity over the prior two weeks across multiple phenotypic dimensions (tic number, frequency, intensity, complexity, interference), some of which are less likely to be directly related to motor cortex functioning. The YGTSS is also a subjective scale based on self-/parent-report and clinician observation that does not always correspond to more objective measures such as video-based tic coding [31]. Future research should continue to examine whether RMT or other TMS-measured neurophysiology indices correspond to tic phenomenology as measured with other questionnaires and/or direct observation measures.

One important limitation within this work is that, although we are inferring developmental information from this dataset, it is a cross-sectional study. Longitudinal research is needed to understand the relationship between changes in brain network maturation, RMT, and tic severity over time.

In conclusion, the current study suggests age-related differences in cortical excitability between individuals with and without TS, likely reflecting a developmental delay in the maturation of functional brain networks in individuals with TS that may normalise with age.

Continued investigation of the developmental trajectory of cortical excitability in TS may help further our understanding of individual differences in disorder course (i.e., distinguish those whose tics do vs. do not improve with age) and reveal opportunities for intervention, particularly with neuromodulatory approaches that directly target neural circuit function.

## Acknowledgements

SRJ, KD, CC, VB, YW, DG, JP, JR and KB would like to acknowledge MRC partnership award funding [MR/Z5050555/1]. KJB reports research funding to his institution from Emalex Biosciences and from Zhittya Genesis Medicine. SRJ is supported by research grants from Medical Research Council (T032588) and (UKRI 527), Parkinson’s UK, and the NIHR Nottingham Biomedical Research Centre. CC is supported by National Institute of Mental Health (R61MH123754), University of Minnesota MnDRIVE (Minnesota’s Discovery, Re-search and Innovation Economy) initiative. YW is supported by National Research Agency (ANR-18-CE37-0008-01) and the program “Investissements d’avenir” ANR-10-IAIHU-0006. DLG was supported by research grants from the US NIH (R01 NS096207). He also reports re-search funding to his institution from Emalex Biosciences, Inc. (clinical trial, Tourette Syndrome), PTC Therapeutics (registry and clinical trial, Amino Acid Decarboxylase Deficiency), Neurocrine Biosciences (clinical trial, cerebral palsy), and Quince Therapeutics (clinical trial, ataxia telangiectasia). He has served as a paid consultant for PTC Therapeutics, Noema Pharma, Acadia Pharmaceuticals, Inc., and Emalex Biosciences and has received travel support from PTC Therapeutics and Emalex Biosciences to attend investigator or scientific meetings. He has provided educational lectures for Illumina, Inc, Emalex Biosciences, and PTC Therapeutics, Inc. He has received book/publication royalties from Elsevier and Wolters Kluwer.

We would like to thank all the participants who took part in the studies and the researchers who assisted with data collection from the original studies. Including: Isabel Farr, University of Nottingham, UK; Aikaterini Gialopsou, University of Nottingham; Hilmar Sigurdsson, Institute of Neuroscience CSIC-UMH, Alicant; Maris Sartoris Therese, Paris Brain Institute - ICM, Sorbonne University, INSERM, CNRS Paris, France; Charles Lamy Jean, Paris Brain Institute - ICM, Sorbonne University, INSERM, CNRS Paris, France; Cyril Atkinson Clement, Sorbonne University, INSERM, CNRS Paris, France; Jennifer Tübing, Institute of Systems Motor Science, University of Lübeck, Lübeck, Germany, Eva Nießen, Cognitive Neuroscience, Institute of Neuroscience & Medicine (INM-3), Research Centre Jülich, Jülich, Germany; Bettina Gigla, Institute of Systems Motor Science, University of Lübeck, Lübeck, Germany; Mo Chen, Department of Psychiatry & Behavioral Sciences, University of Minnesota, Gillette Children’s Hospital, Minnesota, USA, Alana Lieske, Department of Psychiatry & Behavioural Sciences, Masonic Institute for the Developing Brain, University of Minnesota, USA, Suma Jacob, UCLA Semel Institute for Neuroscience and Human Behaviour, Los Angeles, USA

## Supplementary material

**Figure S1:**
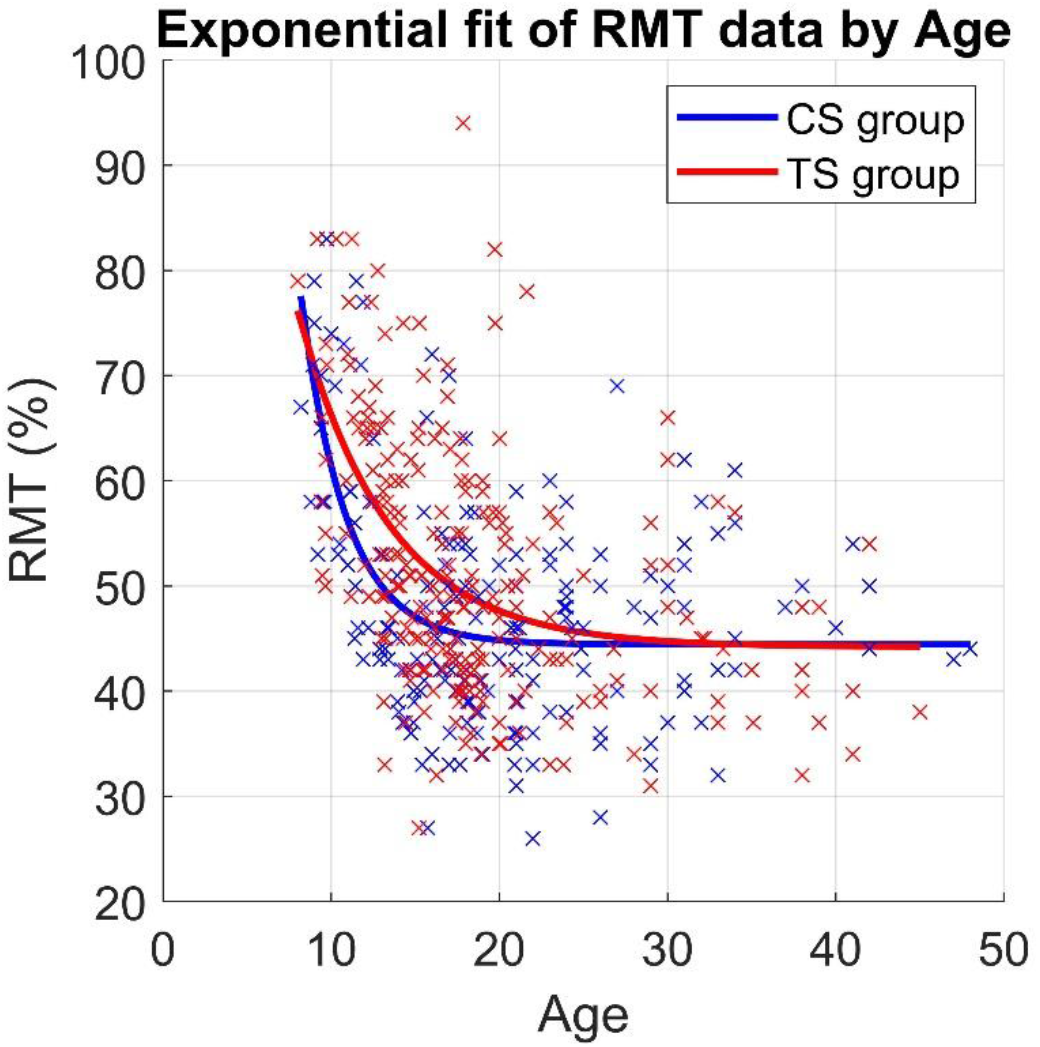
Scatter plot showing individual RMT values as a function of age for the CS group (blue) and the TS group (red) when maximum age is restricted to 50 years. The data from each group was fitted using an exponential function. The solid blue line indicates the fitted function for the CS group and the red line for the TS group. Note both groups show a decrease in RMT with age during adolescence before reaching asymptote as adults.

### Effects of including a small number of older adults

To ensure that the data fitting process was not influenced by including a small or unequal number of older adults, we re-ran the analysis including only participants aged under 50 years (see Figure S1). This analysis confirmed the analyses reported above. Specifically, the analyses confirmed that the observed between-group difference in the rate of decrease in RMT with age during adolescence (k) was statistically significant (observed difference = 0.19, p = 0.005) and that the observed difference in the age at which each group attained adult values of RMT (asymptote) also remained statistically significant (CS asymptote reached at 12.4 years, TS asymptote reached at 24.8 years; p = 0.01). Once again, an independent groups t-test confirmed that once RMT values had plateaued at adult (asymptote) levels RMT they did not differ between the groups (Means: CS group 45.2 ± 8.5 years, TS group 44.5 ± 8.6 years; d = 0.07, t(190) = 0.7).

### Effects of including a biphasic study without control group

One of the studies contributing to the pooled data was conducted using a TMS system that uses biphasic rather than monophasic pulses. TMS pulse type is known to significantly impact RMT [32], and given that this study also only contributed TS data we wanted to ensure our findings remain true with data from this site excluded (N= 48 TS participants). The analysis above was re-run using only data from the ten sites using monophasic TMS pulses. It should be noted that this analysis is included for completeness. Biphasic pulses are known to reduce RMT which would converge from the pattern of results we see whereby individuals with TS have slightly higher RMTs on average with a decreased rate of reduction with age in the TS group.

### Excluding biphasic study without control group – Curve fitting

An identical curve-fitting analyses was conducted with the biphasic study excluded (see Figure S1). The analyses confirmed that the observed between-group difference in the rate of decrease in RMT with age during adolescence (k) parameter was statistically significant (observed difference = 0.13, p = 0.05) and that the observed difference in the age at which each group attained adult values of RMT (asymptote) also remained statistically significant (CS asymptote reached at 12.4 years, TS asymptote reached at 18.7 years; p = 0.05). An independent groups t-test confirmed that once RMT values had plateaued at adult (asymptote) levels RMT did not differ between the groups (Means: CS group 45.2 ± 8.5 years, TS group 44.8 ± 8.2 years; d < −0.0001, t(208) = 0.995).

**Figure S2:**
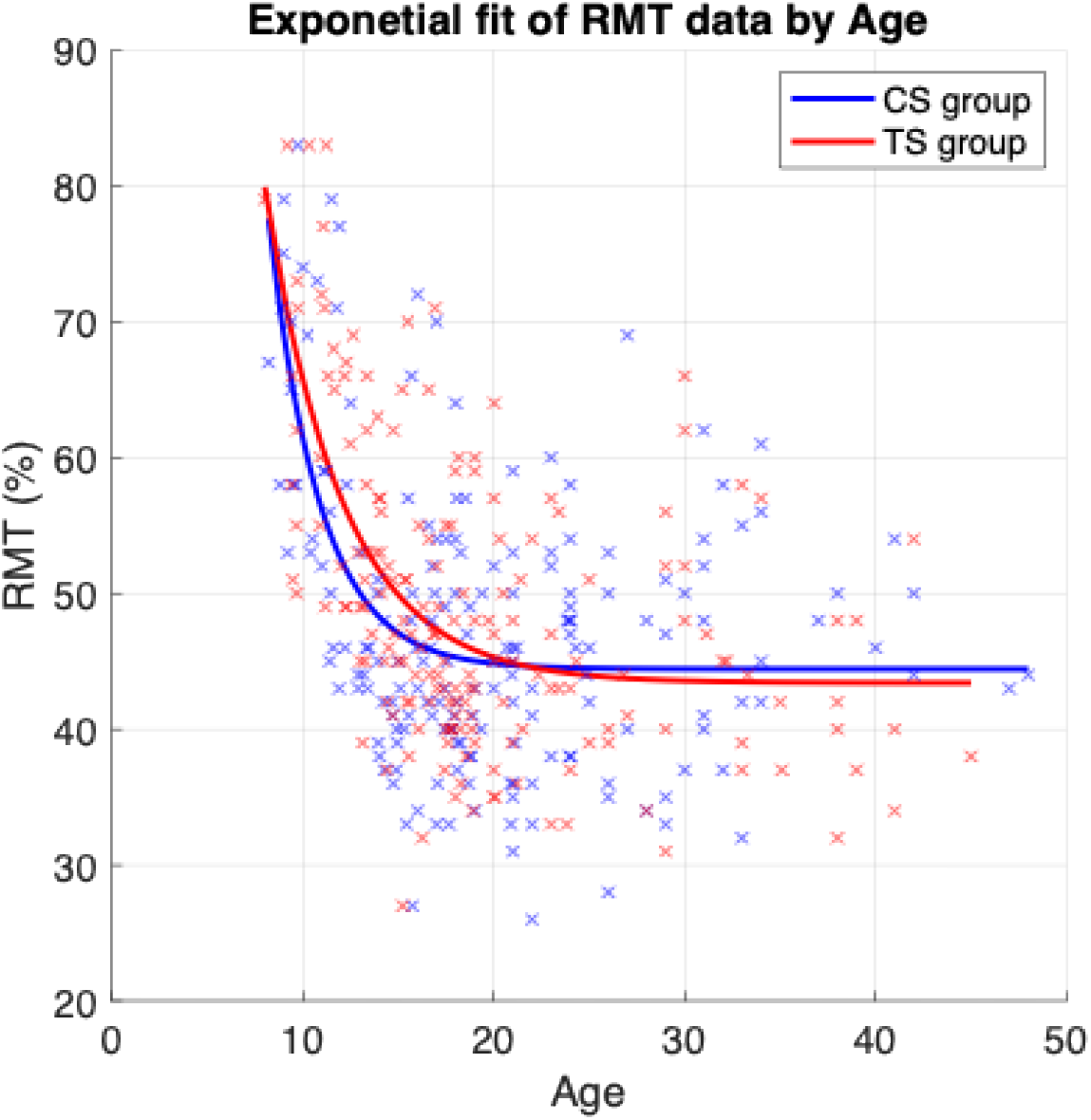
Scatter plot showing individual RMT values as a function of age for the CS group (blue) and the TS group (red) for studies using monophasic TMS pulses only. The data from each group was fitted using an exponential function. The solid blue line indicates the fitted function for the CS group and the red line for the TS group. Note both groups show a decrease in RMT with age during adolescence before reaching an asymptote as adults.

